# Characterizing Human KIF1Bß Motor Activity by Single-Molecule Motility Assays and *Caenorhabtidis elegans* Genetics

**DOI:** 10.1101/2023.11.12.566784

**Authors:** Rei Iguchi, Tomoki Kita, Taisei Watanabe, Kyoko Chiba, Shinsuke Niwa

## Abstract

The axonal transport of synaptic vesicle precursors relies on KIF1A and UNC-104 ortholog motors. In mammals, KIF1Bß is also responsible for the axonal transport of synaptic vesicle precursors. Mutations in KIF1A and KIF1Bß lead to a wide range of neuropathies. While previous studies have revealed the biochemical, biophysical and cell biological properties of KIF1A, and its defects in neurological disorders, the fundamental properties of KIF1Bß remain elusive. In this study, we determined the motile parameters of KIF1Bß through single-molecule motility assays. Additionally, we established simple methods for testing the axonal transport activity of human KIF1Bß using *Caenorhabditis elegans* genetics. Taking advantage of these methods, we demonstrated that these assays enable the detection of reduced KIF1Bß activities both in vitro and in vivo, that is caused by a disease-associated mutation.

## Introduction

Neurons have long neuronal processes called axons (Luo, 2020). The morphology and function of an axon depend on the function of motor proteins that transport cargo organelles by moving in a processive and directional manner along microtubules (Hirokawa et al., 2009). The Kinesin superfamily proteins (KIFs) and cytoplasmic dynein serve as molecular motors for anterograde and retrograde axonal transport, respectively (Holzbaur and Scherer, 2011). Each KIF binds to a specific organelle or protein complex and transports them toward the axon terminal (Hirokawa et al., 2009).

Synaptic vesicles that contain neurotransmitters are essential for synaptic functions (Luo, 2020). Mature synaptic vesicles are generated at the synaptic terminal. However, the components of synaptic vesicles are primarily synthesized in the cell body and then transported to synapses along the axon (Chiba et al., 2023; Hirokawa et al., 2009). The vesicular organelle responsible for this axonal transport, known as a synaptic vesicle precursor, is carried by KIF1A and UNC-104 orthologs in all animals analyzed so far (Chiba et al., 2023). These motor proteins belong to the kinesin-3 family (Hirokawa et al., 2009). UNC-104, a *Caenorhabditis elegans* (*C.elegans*) kinesin, is the founding member of kinesin-3 family (Hall and Hedgecock, 1991; Otsuka et al., 1991). Mutations in *unc-104* gene induces mislocalization of synapses (Hall and Hedgecock, 1991; Otsuka et al., 1991). Mammals have two orthologs of UNC-104 motor in the genome, known as KIF1A and KIF1Bß (Niwa et al., 2008; Okada et al., 1995; Zhao et al., 2001). KIF1A has been identified as a molecular motor responsible for transporting synaptic vesicle precursors in mice (Okada et al., 1995). The genomic locus of KIF1B generates two isoforms of motor proteins, KIF1Bα and KIF1Bß(Zhao et al., 2001). Previous studies have demonstrated that KIF1Bα is involved in the transport of mitochondria, whereas KIF1Bß, structurally similar to KIF1A, also plays a role in transporting synaptic vesicle precursors (Nangaku et al., 1994; Niwa et al., 2008; Zhao et al., 2001). In addition to the axonal transport of synaptic vesicle precursors, studies in invertebrates suggest that KIF1A and UNC-104 orthologs transport active zone precursors and mature synaptic vesicles (Chiba et al., 2023; Pack-Chung et al., 2007). Furthermore, KIF1A and KIF1Bß transport other neuronal cargos such as dense-core vesicles and IGF1 receptor containing vesicles in the axon(Stucchi et al., 2018; Xu et al., 2018). KIF1Bß, together with KIF5, is also implicated in lysosome transport in non-neuronal cells (Guardia et al., 2016). Consistent with this function in non-neuronal cells, KIF1Bß exhibits ubiquitous expression, while KIF1A is dominantly expressed in neurons (Okada et al., 1995; Zhao et al., 2001).

Due to the importance of KIFs in axonal morphology and function, mutations in KIFs are often associated with human neuropathies (Baron et al., 2022; Boyle et al., 2021; Budaitis et al., 2021; Ebbing et al., 2008; Esmaeeli Nieh et al., 2015; Hirokawa et al., 2010; Holzbaur and Scherer, 2011; Klebe et al., 2012; Nakano et al., 2022; Pant et al., 2022; Zhao et al., 2001). Mutations in human KIF1A and human KIF1Bß lead to neuronal diseases (Budaitis et al., 2021; Morikawa et al., 2022; Xu et al., 2018; Zhao et al., 2001). The genomic locus of KIF1A tends to mutate at a relatively higher frequency, and genetic variants in KIF1A cause congenital genetic disorders, called KIF1A-associated neurological disorder (KAND) (Boyle et al., 2021; Chiba et al., 2023). Mutations in KIF1Bß are associated with a peripheral neuropathy known as Charcot-Marie-Tooth type 2A type 1(CMT2A1), probably due to reduced axonal transport in patient neurons (Zhao et al., 2001). KIF1Bß is also identified as a candidate 1p36 tumor suppressor that regulates apoptosis in sympathetic neurons (Munirajan et al., 2008). One possible mechanism of tumor suppression is that KIF1Bß regulates calcineurin activity and mitochondrial dynamics (Li et al., 2016).

Numerous studies have revealed the biochemical properties of both wild type KIF1A and disease-associated KIF1A(Anazawa et al., 2022; Boyle et al., 2021; Chiba et al., 2019; Esmaeeli Nieh et al., 2015; Guardia et al., 2016; Kita et al., 2023b; Lam et al., 2021). The phenotypes of loss of *unc-104* function mutant worms can be rescued by the expression of human KIF1A (Chiba et al., 2019). Taking advantage of these systems, we have developed simple methods to determine the axonal transport capability of disease-associated KIF1A (Lam et al., 2021). In contrast, only a study has analyzed the ATPase activity of KIF1Bß (Zhao et al., 2001). Biophysical motile parameters of KIF1Bß have not been determined yet. There is a need for the development of straightforward and rapid genetic methods to assess the transport activity of disease-associated KIF1Bß. In this study, we have established genetic methods to determine whether a mutation in KIF1Bß results in a loss of transport activity. Furthermore, we have measured the motile parameters of KIF1Bß in vitro.

## Results

### *unc-104* mutant worms can be rescued by the expression of human KIF1Bß

KIF1Bß, along with KIF1A, is an mammalian ortholog of UNC-104 motor and has a very similar domain architecture (Fig 1A) (Hirokawa et al., 2009; Niwa et al., 2008). We have previously demonstrated that the expression of human KIF1A can rescue the body movement defects in *unc-104(e1265)*, a widely used loss-of-function allele of *unc-104* (Chiba et al., 2019; Hall and Hedgecock, 1991). In this paper, we refer to this allele as *unc-104(lf)*. Subsequently, we conducted a similar experiment using human KIF1Bß (Niwa et al., 2008). Human KIF1Bß cDNA was fused with the *unc-104* promoter, a neuron specific promoter (Chiba et al., 2019), and expressed in *unc-104(lf)*. Our finding revealed that the expression of human KIF1Bß in worm neurons could rescue the body movement defects observed in the *unc-104(lf)* allele (Fig 1B-D). We have previously shown that KIF1A with disease-associated loss-of-function mutations cannot rescue *unc-104(lf)* (Chiba et al., 2019). To determine if we can detect disease-associate defects in KIF1Bß, we introduced a CMT2A1-associated Q98L mutation into KIF1Bß cDNA and tested whether this mutant version of KIF1Bß could rescue the *unc-104(lf)* mutant or not. Our results indicated that KIF1Bß(Q98L) was unable to rescue the body movement defects (Figs 1E and F).

**Figure 1.**
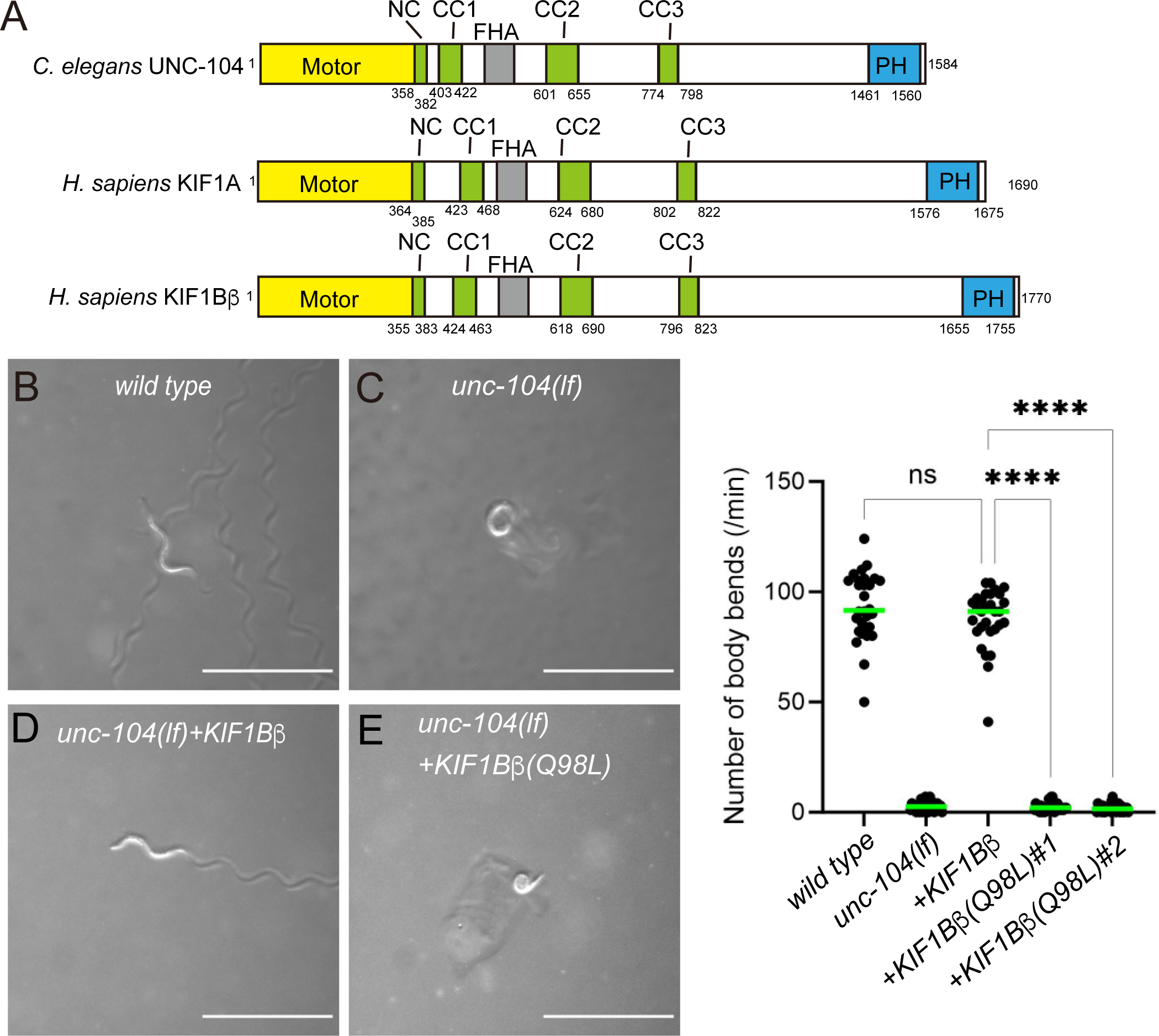
Worm complementation assays. (A) The domain structure of *C. elegans* UNC-104, human KIF1A and human KIF1Bβ proteins. Motor, motor domain. NC, neck coiled-coil domain. CC1, Coiled-coil 1 domain. FHA, Forkhead-associated domain. CC2, Coiled-coil 2 domain. CC3, Coiled-coil 3 domain. PH, Pleckstrin-homology domain. The numbers indicates amino acid numbers. (B-E) Macroscopic phenotypes of *wild type* (B), *unc-104(lf)* (C), *unc-104(lf)* expressing human KIF1Bβ (D) and *unc-104(lf)* expressing KIF1Bβ(Q98L). Scale bars: 1 mm. Note that when wild type KIF1Bβ, but not KIF1Bβ(Q94L), was expressed, motility of *unc-104(lf)* was recovered. (F) Dot plots showing the results of the swimming assay. The number of body bends in a water drop was counted for 1 minute and plotted. Dots represent the data points, green bars represent median value. n = 30 worms. Ordinary one-way ANOVA followed by Tukey’s multiple comparison test. ****, adjusted P Value < 0.0001.

### Human KIF1Bß rescues synaptic defects in *unc-104(lf)*

We proceeded to analyze whether the expression of human KIF1Bß could rescue the synaptic defects in *unc-104(lf)* worms. DA9 neuron is a polarized neuron with distinct regions, including a dendrite, cell body and axon (Fig 2A and B) (Klassen and Shen, 2007). En passant synapses are formed along the axon of the DA9 neuron, making it a valuable model for studying axonal transport and synaptogenesis (Balseiro-Gomez et al., 2022; Glomb et al., 2023; Higashida and Niwa, 2023; Klassen and Shen, 2007; Klassen et al., 2010; Niwa et al., 2016; Wu et al., 2013). Previous studies have shown that synaptic vesicles, visualized by GFP::RAB-3, are mislocalized in the dendrite and cell body of DA9 neuron in *unc-104(lf)* worms while synaptic vesicles are localized to dorsal axon in wild-type DA9 neuron(Niwa et al., 2016; Wu et al., 2013) (Fig 2B and C). When human KIF1Bß was expressed in *unc-104(lf)* mutant worms using the *unc-104* promoter, the localization of synaptic vesicles along the axon in transgenic strains were restored and we could not distinguish them from the *wild-type* neuron (Fig 2D). However, when human KIF1Bß(Q98L) was expressed in *unc-104(lf)* mutants, synaptic vesicles remained mislocalized along the dendrite (Fig 2E and F). We conducted statistical analyses by counting the number of GFP::RAB-3 puncta in the axon and the commissure and dendritic region (Figs 2 G and H). The expression of wild-type KIF1Bß, but not KIF1Bß(Q98L), restored the number of GFP::RAB-3 puncta to the wild-type level in the axon (Fig 2G). Similarly in the commissure and dendrite, the expression of wild-type KIF1Bß, but not KIF1Bß(Q98L), reduced the number of mislocalized GFP::RAB-3 puncta (Fig 2H). The results from these assays suggest that human KIF1Bß is capable of performing axonal transport in the worm neuron, and this assay is useful to detect reduced transport activity caused by disease-associated KIF1Bß mutations.

**Figure 2.**
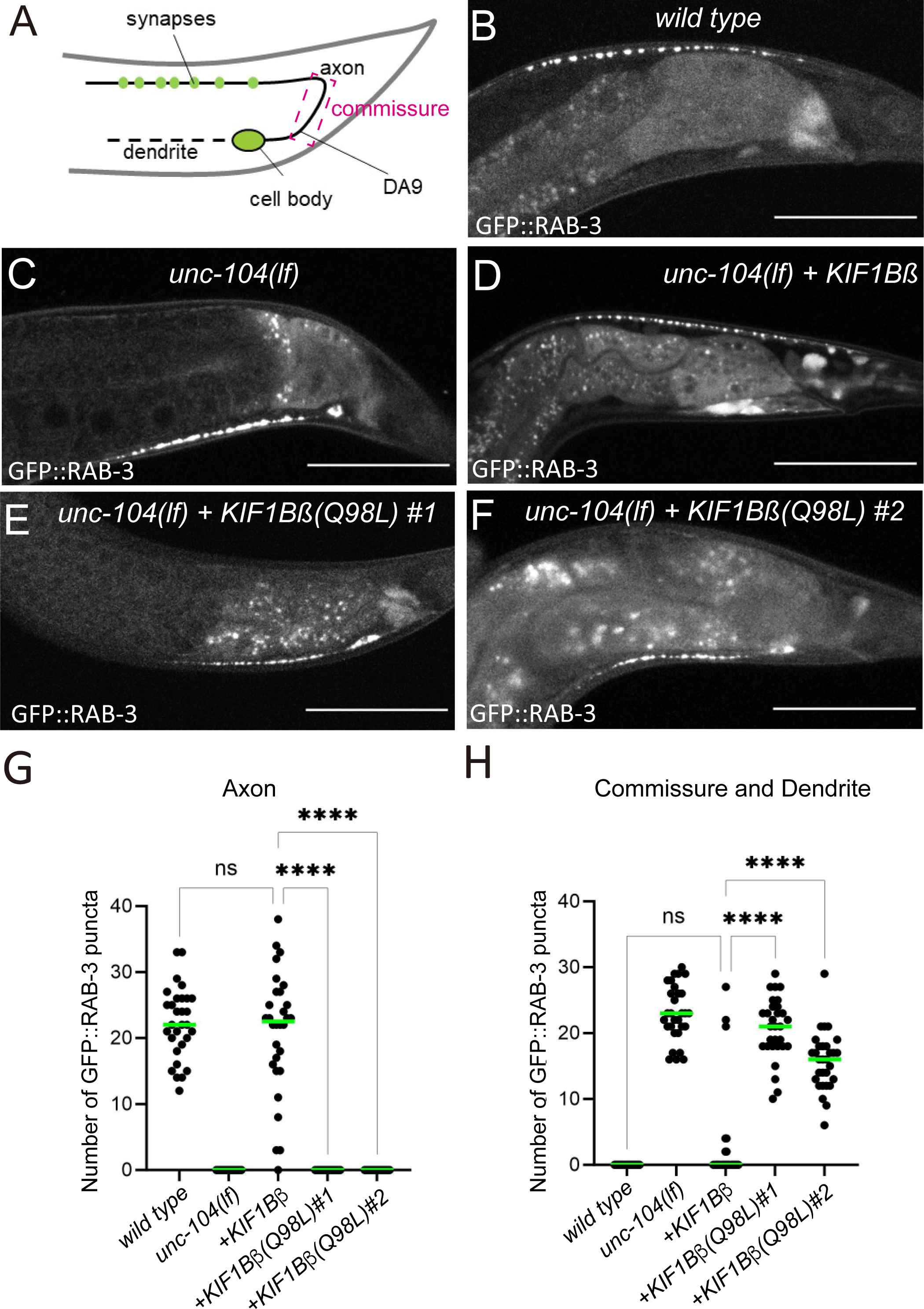
(A) Schematic drawing shows the morphology of the DA9 neuron. Green dots show the localization of the synaptic vesicles or their precursors represented by GFP::RAB-3. (B-F) GFP::RAB-3 was expressed in DA9 neuron under the *itr-1* promoter. Representative images of the localization of GFP∷RAB-3 in *wild type* (B), *unc-104(lf)* (C), *unc-104(lf)* expressing wild type KIF1Bβ (D), *unc-104(lf)* expressing KIF1Bβ(Q98L) (E and F). Bars, 50 µm. (G) Dot plots showing the number of GFP::RAB-3 puncta in the axon of DA9. Dots represent the number of GFP::RAB-3 puncta in each worm, green bars represent median value. n = 30 worms. Ordinary one-way ANOVA followed by Tukey’s multiple comparison test. ****, adjusted P Value < 0.0001. (H) Dot plots showing the number of GFP::RAB-3 puncta in the commissure and the dendrite of DA9. Dots represent the number of GFP::RAB-3 puncta in each worm, green bars represent median value. n = 30 worms. Ordinary one-way ANOVA followed by Tukey’s multiple comparison test. ****, adjusted P Value < 0.0001.

### Human KIF1Bß is a processive motor protein

Detailed physical parameters of KIF1Bß in vitro have remained largely elusive. This contrasts with the case of KIF1A, where a number of studies have revealed physical parameters. Therefore, we measured the motile parameters of KIF1Bß using TIRF assay. We expressed and purified recombinant human KIF1Bß fused with a fluorescent protein sfGFP (KIF1Bß-FL) using Sf9 insect cells and baculovirus. Additionally, we analyzed a deletion mutant of KIF1Bß fused with sfGFP, which consists of the motor, neck coil, FHA, CC1 and CC2 domains (KIF1Bß(1-721)) (Fig 3A and Supplementary Figure S1), as similar deletion mutants of human KIF1A and worm UNC-104 have been analyzed in previous studies (Hammond et al., 2009; Kita et al., 2023a; Tomishige et al., 2002). Both KIF1Bß-FL and KIF1Bß(1-721) showed processive movement on microtubules (Figs 3B and C). The velocity of KIF1Bß-FL and KIF1Bß(1-721) was 0.82 ± 0.44 µm/sec and 0.79 ± 0.34 µm/sec, respectively (Fig 3D, n=342 and 425 molecules, with no statistical difference in t-test). The run length of KIF1Bß-FL and KIF1Bß(1-721) was 7.7 ± 6.7 µm and 8.0 ± 6.7 µm (Fig 3E, n = 342 and 425 molecules, with no significant difference in Mann-Whitney test). These results indicate that the properties of the motor domain were not affected by the deletion of the tail domain. In contrast, the landing rate showed a significant difference. The landing rate of KIF1Bß-FL and KIF1Bß(1-721) was 0.0016 ± 0.0006 µm^-1^sec^-1^ at 2 nM and 0.0062 ± 0.0023 µm^-1^sec^-1^ at 0.5 nM, respectively (Fig 3F). The concentrations could not be matched because, when one was adjusted to an ideal concentration for measurement, the other either became saturated or fell below the limit concentration. Although the concentration of KIF1Bß(1-721) was lower than KIF1Bß-FL in the measurement, the landing rate of KIF1Bß(1-721) was higher (n = 31 and 34 microtubules, p < 0.0001, Mann-Whitney test). This result suggests that KIF1Bß(1-721) is capable of binding to microtubules much more frequently than KIF1Bß-FL, indicating that the domain encoding KIF1Bß(722-1770) inhibits the association of KIF1Bß-FL with microtubules. This is consistent with the predicted KIF1Bß structure obtained by the Alphafold2 (Fig S1) as described in the Discussion.

**Figure 3.**
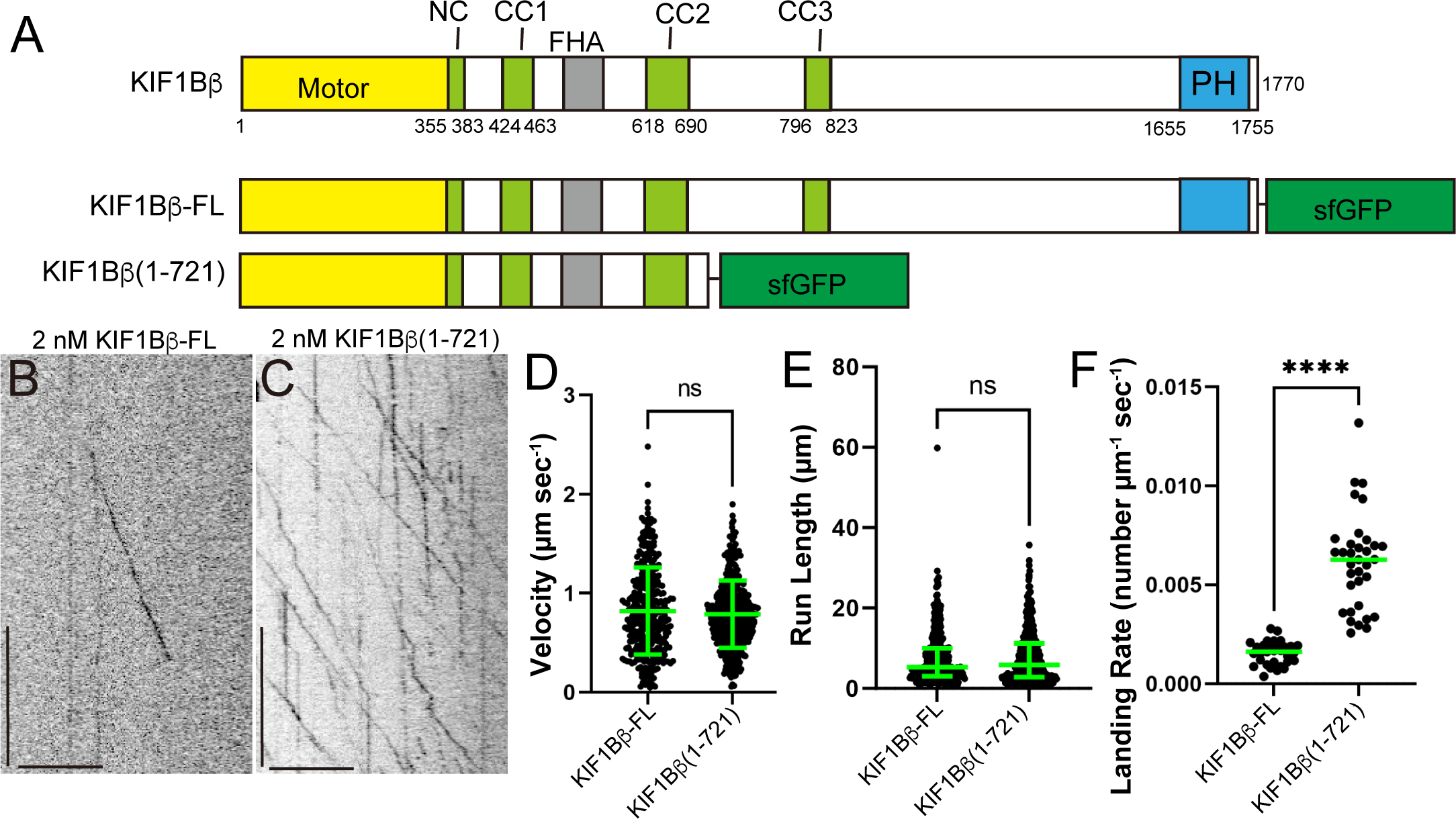
Single molecule motility assays. (A) Schematic drawing of the domain organization of human KIF1Bß motor protein and the full length of KIF1Bß fused with sfGFP (KIF1Bß-FL) and a deletion mutant of KIF1Bß, consisting of the motor domain to CC2 domain, fused with sfGFP (KIF1Bß(1-721)). (B and C) Representative kymographs showing the motility of 2 nM KIF1Bß-FL dimer and KIF1Bß(1-721) dimer in the presence of 2 mM ATP. Vertical and horizontal bars show 10 seconds and 10 µm, respectively. (D) Dot plots showing the velocity of KIF1Bß-FL and KIF1Bß(1-721). Each dot shows a velocity of a molecule. Green bars represent mean ± S.D.. n = 342 and 425 for KIF1Bß-FL and KIF1Bß(1-721), respectively. ns, no statistically significant difference in t-test. (E) Dot plots showing the run length of KIF1Bß-FL and KIF1Bß(1-721). The distance each molecule traveled to bind to and leave the microtubule was measured. Each dot shows a run length of a molecule. Green bars represent median value and interquartile range. n = 342 and 425 for KIF1Bß-FL and KIF1Bß(1-721), respectively. ns, no statistically significant difference in Mann-Whitney U test. (F) Dot plots showing the landing rate of 2 nM KIF1Bß-FL and 0.5 nM KIF1Bß(1-721). The number of molecules bound to a 1-µm region of microtubules per second is shown. Each dot shows a single datum point. Green bars represent median value. n = 31 and 34 microtubules for KIF1Bß-FL and KIF1Bß(1-721), respectively. Mann-Whitney U test. ****, p < 0.0001. Note that the concentrations of the two measurements are different because of the different concentrations suitable for counding molecules.

### KIF1Bß(Q98L) reduces the motor activity

It has been demonstrated that physical parameters affected by a disease-associated mutation correlate with human symptoms in the case of KIF1A(Boyle et al., 2021). In contrast in the case of KIF1Bß, Q98L mutation, which has been found in a Japanese CMT2A family, was studied by the ATPase assay only (Zhao et al., 2001). Therefore, we conducted a comparison between the activity of wild-type KIF1Bß(1-721) (KIF1Bß(1-721)wt) and KIF1Bß(1-721)(Q98L) using single molecule assays. In contrast to KIF1Bß(1-721)wt, which exhibits processive movement on microtubules, KIF1Bß(1-721)(Q98L) displayed slower movement on microtubules (Fig 4A-C). The run length of KIF1Bß(1-721)(Q98L) was shorter than KIF1Bß (1-721)wt (Fig 4D). Moreover, the landing rate of KIF1Bß(1-721)(Q98L) was significantly reduced compared to KIF1Bß (1-721)wt (Fig. 4E). Collectively, these data suggest that the KIF1Bß(Q98L) mutation reduces a broad range of motile parameters in the human KIF1Bß motor protein.

**Figure 4.**
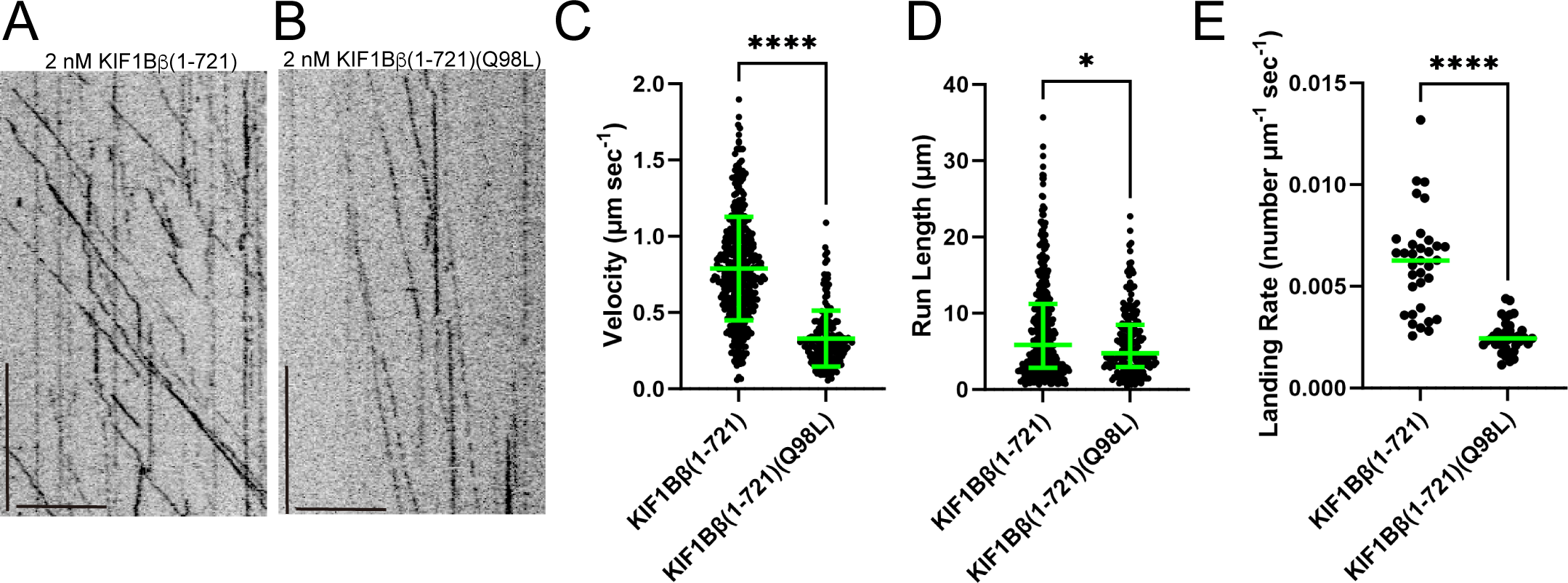
Analysis of the Q98L mutation. (A and B) Representative kymographs showing the motility of 2 nM KIF1Bß(1-721) dimer and KIF1Bß(1-721)(Q98L) dimer in the presence of 2 mM ATP. Vertical and horizontal bars show 10 seconds and 10 µm, respectively. (C) Dot plots showing the velocity of KIF1Bß(1-721) and KIF1Bß(1-721)(Q98L). Each dot shows the velocity of each molecule. Green bars represent mean ± S.D.. n = 425 and 208 for KIF1Bß(1-721) and KIF1Bß(1-721)(Q98L), respectively. t-test. ****, p < 0.0001. KIF1Bß(1-721) values are replotted from Fig. 3D. (D) Dot plots showing the run length of KIF1Bß(1-721) and KIF1Bß(1-721)(Q98L). Each dot shows a single datum point. Green bars represent median value and interquartile range. n = 425 and 208 for KIF1Bß(1-721) and KIF1Bß(1-721)(Q98L), respectively. Mann-Whitney U test. *, p = 0.0465. KIF1Bß(1-721) values are replotted from Fig. 3E. (E) Dot plots showing the landing rate of KIF1Bß(1-721) and KIF1Bß(1-721)(Q98L). Each dot shows a single datum point. Green bars represent median value. n = 34 and 37 microtubules for KIF1Bß(1-721) and KIF1Bß(1-721)(Q98L), respectively. Mann-Whitney U test. ****, p < 0.0001. KIF1Bß(1-721) values are replotted from Fig. 3F.

### Establishment of unc-104(Q94L) worms by genome editing

The residue KIF1Bß(Q98L) is well conserved in worm UNC-104 and is equivalent to UNC-104(Q94L) (Figure 5A). We next introduced this mutation into *C. elegans* using CRISPR/cas9 (Arribere et al., 2014; Ghanta et al., 2021). Introduction of the mutation was confirmed by genomic PCR followed by a restriction enzyme digestion and Sanger sequencing (Fig S2). We then observed the macroscopic phenotypes of homozygous worms with Q94L mutation (Fig 5B and C). Unlike strong loss-of-function alleles of *unc-104* (Fig 1C) (Anazawa et al., 2022), *unc-104(Q94L)* mutant worms exhibited movement on feeder plates (Fig 5B and C). Unlike KAND model *unc-104* mutants (Anazawa et al., 2022), the body size of the mutant worms was comparable to that of wild-type worms (Fig 5B and C). However, the movement of mutant worms was slower than that of wild-type worms. To quantitively analyze the worm movement, we counted the number of body bends in a water drop (Fig 5D). The assay confirmed that the body movement was 30% less frequent than wild type (Fig 5D). Next, we observed the synaptic vesicle marker RAB-3 in the DA9 neuron of *unc-104(Q94L)* worms (Figs 5E-G). We counted the number of GFP::RAB-3 puncta in the commissure and dendrite (Fig 5G). Because no accumulation was observed in the wild type background, we consider the number to reflect the severity of the phenotype. While the accumulation of GFP::RAB-3 was not observed in the commissure and dendrite in wild type worms, 23% of *unc-104(Q94L)* mutant worms exhibited mis-accumulation of GFP::RAB-3 in the commissure and dendrite (Fig 5G). These data suggests that unc-104(Q94L) is a hypomorphic allele of *unc-104*.

**Figure 5.**
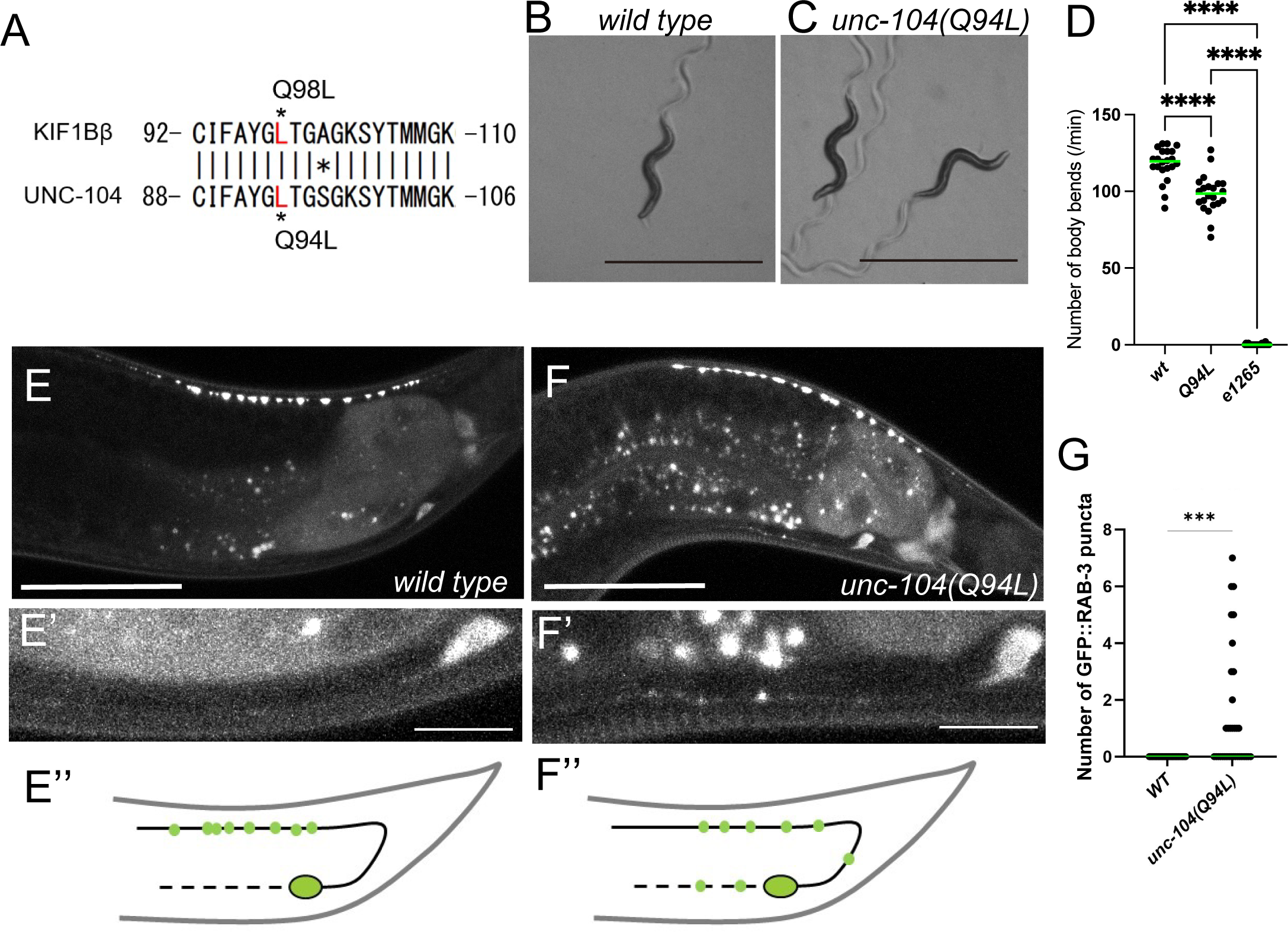
Establishment of CMT2A type 1 model worms. (A) Sequence comparison between human KIF1Bß and *C. elegans* UNC-104. (B and C) Macroscopic phenotypes of *wild type* (B) and unc-104(Q94L) homozygotes (C) at 1-day adults. Bars, 1 mm. (D) Dot plots showing the results of the swimming assay. The number of body bends in a water drop was counted for 1 minute and plotted. Dots represent the data points and green bars represent median value. n = 22 worms. Ordinary one-way ANOVA followed by Tukey’s multiple comparison test. ****, adjusted P Value < 0.0001. (E and F) GFP::RAB-3 was expressed in DA9 neuron under the *itr-1* promoter. Representative images of the localization of GFP∷RAB-3 in *wild type* (E), *unc-104(Q94L)* (F). Bars, 50 µm. (E’) and (F’) show magnified images of the dendritic region of panel (E) and (F). Bars, 10 µm. (E’’) and (F’’) show schematic drawings of the localization of GFP::RAB-3. (G) Dot plots showing the number of GFP::RAB-3 puncta in the commissure and the dendrite of DA9. Each dot represents the number of GFP::RAB-3 puncta in each worm. Ordinary one-way ANOVA followed by Tukey’s multiple comparison test. n = 30 and 34 in *wild type* and *unc-104(Q94L)* worms, respectively. ***, adjusted P Value < 0.001.

### Analysis of *unc-104(Q94L)/unc-104(lf)* transheterozygotes

We have previously observed weak mis-accumulation of GFP::RAB-3 in the dendrite in gain of function alleles of *unc-104* as well (Niwa et al., 2016). To exclude the possibility that *unc-104(Q94L)* might be a gain-of-function allele and to confirm that *unc-104(Q94L)* is a hypomorphic allele, we crossed *unc-104(Q94L)* worms and *unc-104(lf)* to generate transheterozygotes, *unc-104(Q94L)/unc-104(lf)* (Fig 6A-F). As described in Figure 2B, 2C and 5F, the synaptic marker RAB-3 localized in dorsal axon in DA9 neuron of wild type worms (Fig 6A), mislocalized to the dendrite in *unc-104(lf)* worms (Fig 6B), and partically mislocalized to the dendrite and commisure in *unc-104(Q94L)* worms (Fig 6C). When wild type or *unc-104(Q94L)* worms were crossed with *unc-104(lf)* worms, while *wild type/unc-104(lf)* heterozygotes (+/*unc-104(lf)* in the figure) did not exhibit GFP::RAB-3 mislocalization in DA9 neuron, *unc-104(Q94L)/unc-104(lf)* transheterozygotes exhibited misaccumulation of GFP::RAB-3 in the dendrite and commissure (Fig 6D and E). To confirm the phenotype, we counted the number of GFP::RAB-3 puncta in the commissure and dendrite (Fig. 6F). We found the mislocalization of GFP::RAB-3 was increased in *unc-104(Q94L)/unc-104(lf)* transheterozygotes compared to *wild type/unc-104(lf)* heterozygotes and *unc-104(Q94L)* homozygotes. These data again suggests that unc-104(Q94L) is a hypomorphic allele of *unc-104*.

**Figure 6.**
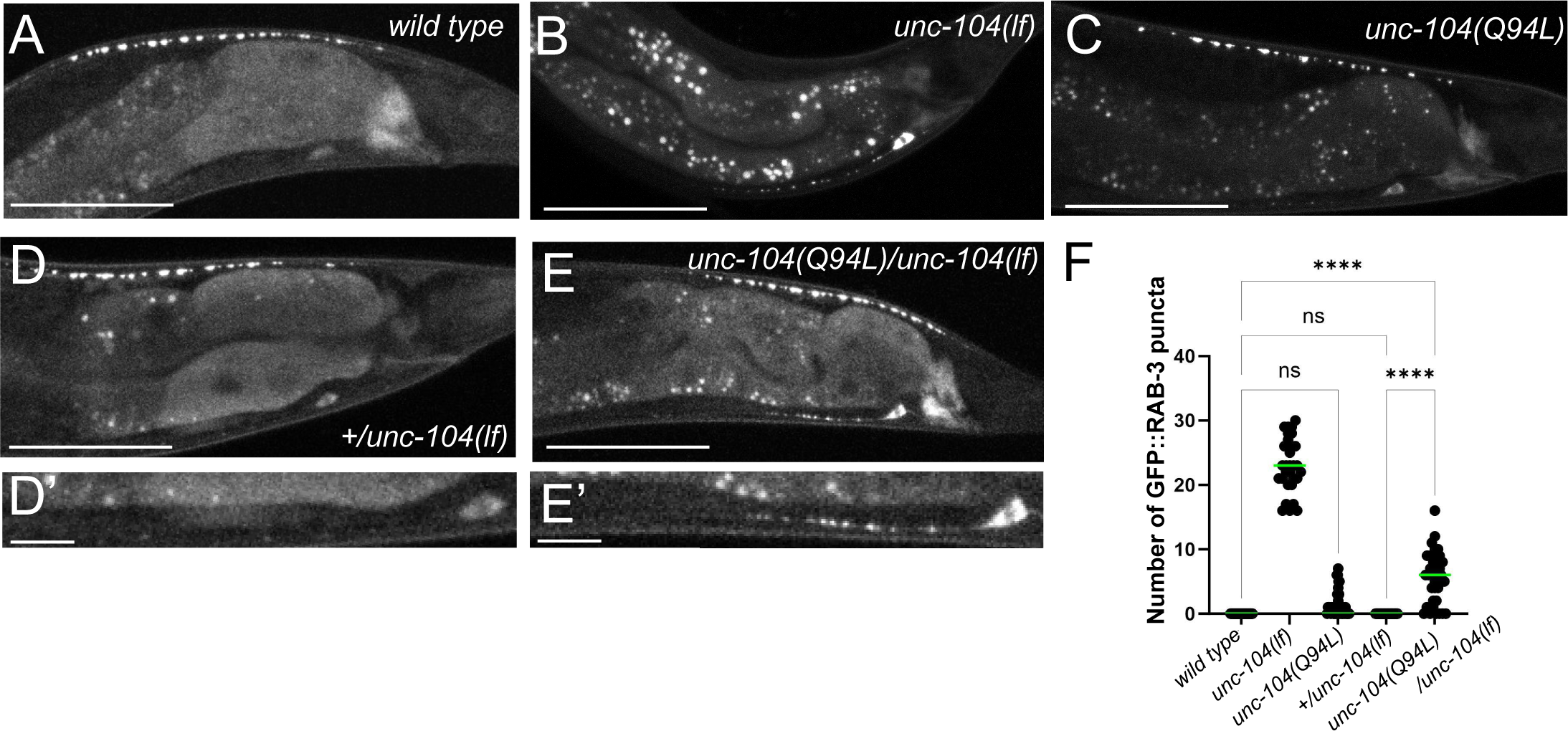
Transheterozygote analysis. (A-E) GFP::RAB-3 was expressed in DA9 neuron under the *itr-1* promoter. Representative images of the localization of GFP∷RAB-3 in *wild type* (A), *unc-104(lf)* (B), *unc-104(Q94L)* (C), *+*/*unc-104(lf)* heterozygote (D), and *unc-104(Q94L)/unc-104(lf)* transheterozygote (E). Bars, 50 µm. (D’) and (E’) show magnified images of the dendritic region of panel (D) and panel (E). Bars, 10 µm. (F) Dot plot showing the number of GFP::RAB-3 puncta in the commissure and the dendrite of DA9. Ordinary one-way ANOVA followed by Tukey’s multiple comparison test. n = 30, 40, 40, 33 and 40 in *wild type*, *unc-104(lf)*, *unc-104(Q94L)*, *+/unc-104(lf)* and *unc-104(Q94L)/unc-104(lf)*, respectively. ****, adjusted P Value < 0.0001.

## Discussion

### Basic properties of human KIF1Bß

The role of both KIF1A and KIF1Bß as axonal motors responsible for the transport of synaptic vesicle precursors in mammals has been shown (Niwa et al., 2008; Okada et al., 1995; Zhao et al., 2001). While KIF1A has been subjected to extensive analysis of its biochemical and biophysical properties, the examination of KIF1Bß has been relatively limited. Although the ATPase activity of KIF1Bß has been measured (Zhao et al., 2001), the biophysical properties of human KIF1Bß have not been determined. In this study, we analyzed biophysical parameters of KIF1Bß using single molecule motility assays. The velocity, run length, and landing rate of KIF1Bß are almost identical to those of KIF1A (Chiba et al., 2019), which is consistent with the redundant functions of these two motors. Similar to KIF1A, KIF1Bß can transport synaptic materials in worm neurons and can rescue defects in *unc-104(lf)* mutant worms (Fig 1A and B).

### Autoinhibition of KIF1Bß

In our single molecule assays conducted in the absence of cargos, we observed that KIF1Bß(1-721) exhibited more frequent microtubule binding compared to KIF1Bß-FL (Fig 3). This property is similar to the behavior of KIF1A as reported previously (Budaitis et al., 2021). While the precise mechanism underlying this phenomena needs further analysis, this would be related to the regulatory mechanism known as autoinhibition, which is a common feature among motor proteins (Verhey and Hammond, 2009). Kinesin motors are generally folded in the absence of cargo molecules and kept in an inactive state. For instance, kinesin-1 is known to adopted a folded conformation in the absence of cargo (Tan et al., 2023; Weijman et al., 2022). A common property of kinesin-3 motors is that the CC1 and CC2 domains fold toward the motor domain and inhibit the activity of the motor domain (Ren et al., 2018; Wang et al., 2022). While the structural details of full-length KIF1Bß orthologs have not been revealed, a recent study has clarified the crystal structure of the full-length KLP-6 (Wang et al., 2022), a kinesin-3 family motor unique to worms. Their study revealed that not only the CC1 and CC2 domains but also more C-terminal tail domains of KLP-6 are folded toward the motor domain, and inhibit the activity of KLP-6 (Wang et al., 2022). In alignment of this finding, it is likely that KIF1Bß(722-1770), the C-terminal region of KIF1Bß containing cargo-binding domains, may inhibit the activity of KIF1Bß. Consistent with this idea, the AlphaFold2 prediction suggests that the region encoded by KIF1Bß(722-1770) adopts a folded conformation toward KIF1Bß(1-721) (Supplementary Figure S1), which is similar to kinesin-1(Tan et al., 2023; Weijman et al., 2022). To fully understand the mechanism of the autoinhibition of KIF1Bß and its orthologs, further studies focusing on elucidating the high-resolution structure of the full-length proteins will be essential.

### KIF1Bß(Q98L) mutation reduces the transport activity

The ongoing revolution in the whole-genome sequence technology is revealing a lot of disease associated mutations in KIF1A and KIF1Bβ (Boyle et al., 2021; Xu et al., 2018). When previously uncharacterized single nucleotide polymorphisms are identified in patients, analyzing these mutations is required. In the case of KIF1A mutations, we and others have established genetic and biochemical assays to test whether a mutation results in a loss of function (Anazawa et al., 2022; Boyle et al., 2021; Budaitis et al., 2021; Chiba et al., 2019; Lam et al., 2021). Using these assays, numerous disease-associated KIF1A mutations have been characterized. Comprehensive biochemical analysis showed that such disease-associated mutations in KIF1A affect various motile parameters of the KIF1A motor (Boyle et al., 2021). In contrast, methods to analyze KIF1Bß mutations were yet to be developed. While it has been reported that the KIF1Bß(Q98L) is associated with CMT2A in a Japanese family, other studies have shown that CMT2A is caused by mutations in MFN2, a protein essential for mitochondrial fusion(Zuchner et al., 2004). The KIF1B locus is in close proximity to the MFN2 locus on the human chromosome 1 (Zuchner et al., 2004). These make it unclear whether or not the KIF1Bß(Q98L) mutation really affects neuronal function or not(Zuchner et al., 2004). Our in vivo and in vitro assays gave compelling evidence demonstrating that the KIF1Bß(Q98L) mutation reduces motor activity and disrupts the axonal transport synaptic vesicle precursors. Thus, it is reasonable to conclude that KIF1Bß(Q98L) mutation induces an adverse impact on neuronal function in human neurons (Zhao et al., 2001). Interestingly, mutations in KIF1Bß have been linked to neuroblastoma (Munirajan et al., 2008). However, whether these mutations affect the transport activity of KIF1Bß has not been revealed. The system we have developed here would be useful to elucidate the molecular mechanism underlying these phenomena and advancing our understanding of KIF1Bß-related pathologies.

## Methods

### Worm experiments

*C. elegans* strains were maintained as described previously (Brenner, 1974). N2 wild-type worms and OP50 feeder bacteria were obtained from the *C. elegans* genetic center (CGC) (Minneapolis, MN, USA). *wyIs85* and *unc-104(e1265)*; *wyIs85* were described previously (Anazawa et al., 2022). Plasmids to transform worms are described in ***supplementary table S1***. Human KIF1Bß cDNA encoding isoform c of KIF1Bß was obtained from Kazusa DNA Institute (Chiba, Japan). We found mutations and insertions in Kazusa’s cDNA clone. Firstly, mutations were corrected by PCR-based mutagenesis (Stratagene). Secondly, insertions in the Kazasa’s clone were deleted by PCR to obtain cDNA encoding KIF1Bß isoform b. cDNA encoding KIF1Bß isoform b was inserted to *Punc-104* endoding vector that is previously described (Chiba et al., 2019). Transformation of *C. elegans* was performed by plasmid injection as described (Mello et al., 1991). Strains used in this study are described in ***supplementary table S2***. The swim test was performed as described previously (Pierce-Shimomura et al., 2008).

### Genome editing

*unc-104(Q94L)* mutation was introduced by Alt-R® CRISPR/cas9 system (Integrated DNA Technologies). Alt-R® CRISPR-Cas9 tracrRNA (#1072533), crRNA for *unc-104* and Alt-R® *S.p.* Cas9 (#1081058) were purchased from Integrated DNA Technologies. The target sequence was 5’-GCATACGGTCAAACAGGATC −3’ (Supplementary Figure S2) The repair template was 5’-AATAATTGACTTCTAGGTATAATGTCTGCATTTTTGCATACGGTCTAACCGGATCCGGAAAATCATATACAATGATGGGAAAAGCCAATG −3’ Injection mix was prepared as described with a slight modification (Ghanta et al., 2021). We excluded *rol-6* marker because successful injections should produce mutants with *unc* phenotypes due to deletion mutations of *unc-104* caused by repair failures. 3-days later after the injection, we selected a few plates that contain strong *unc* worms. Then, *unc* mutant worms as well as superficially *wild-type* worms were singled from these plates. 7 days later, plates that contain unc-104(Q94L) allele was selected by genomic PCR and BamHI (NEB) digestion. 30% of plates contained either *unc-104(Q94L)* heterozygotes or homozygotes. It was revealed that strong unc worms were deletion mutant of *unc-104*.

### Statistical analyses and graph preparation

Statistical analyses were performed using Graph Pad Prism version 9. Statistical methods are described in the figure legends. Graphs were prepared using Graph Pad Prism version 9, exported in the TIFF format and aligned by Adobe Illustrator 2021.

### Purification of recombinant KIF1Bß

Reagents were purchased from Nacarai tesque (Kyoto, Japan), unless described. Plasmids to express recombinant KIF1Bß are described in ***supplementary table S1***. Sf9 cells (Thermo Fisher Scientific) were maintained in Sf900^TM^ II SFM (Thermo Fisher Scientific) at 27°C. DH10Bac (Thermo Fisher Scientific) were used to generate bacmid. To prepare baculovirus, 2 × 10^6^ cells of Sf9 cells were transferred to each well of a 6 well plate (FALCON). After the cells attached to the bottom of the dishes, about ∼5 μg of bacmid were transfected using 5 μL of TransIT^®^-Insect transfection reagent (Takara Bio Inc.). 5 days after initial transfection, the culture media were collected and spun at 3,000× g for 10 min to obtain the supernatant containing recombinant baculovirus (P1). To express recombinant proteins, 200 mL of Sf9 cells (2 × 10^6^ cells/mL) were infected with 100 µL of P1 virus and cultured for 65 h at 27°C. Cells were harvested and stocked at −80°C. Sf9 cells were resuspended in 30 mL of lysis buffer (50 mM HEPES-KOH, pH 7.5, 150 mM KCH3COO, 2 mM MgSO4, 1 mM EGTA, 10% glycerol) along with 1 mM DTT, 1 mM PMSF, 0.1 mM ATP and 0.5% Triton X-100.

After incubating on ice for 10 min, lysates were cleared by ultracentrifugation (80,000 × g, 20 min, 4°C). Lysate was loaded on Streptactin-XT resin (IBA Lifesciences, Göttingen, Germany) (bead volume: 2 ml). The resin was washed with 40 ml wash buffer (50 mM HEPES-KOH, pH 8.0, 450 mM KCH_3_COO, 2 mM MgSO_4_, 1 mM EGTA, 10% glycerol). Protein was eluted with 40 ml elution buffer (50 mM HEPES-KOH, pH 8.0, 150 mM KCH_3_COO, 2 mM MgSO_4_, 1 mM EGTA, 10% glycerol, 100 mM biotin). Eluted fractions were further separated by NGC chromatography system (Bio-Rad) equipped with a Superdex 200 Increase 10/300 GL column (Cytiva). Peak fractions were collected and frozen.

### TIRF single-molecule motility assays

TIRF assays were performed as described (Chiba et al., 2019; Kita et al., 2023a). Glass chambers were prepared by acid washing as previously described (Chiba et al., 2022). Glass chambers were coated with PLL-PEG-biotin (SuSoS, Dübendorf, Switzerland). Polymerized microtubules were flowed into streptavidin adsorbed flow chambers and allowed to adhere for 5–10 min. Unbound microtubules were washed away using assay buffer (90 mM HEPES-KOH pH 7.4, 50 mM KCH_3_COO, 2 mM Mg(CH_3_COO)_2_, 1 mM EGTA, 10% glycerol, 0.1 mg/ml biotin–BSA, 0.2 mg/ml kappa-casein, 0.5% Pluronic F127, 2 mM ATP, and an oxygen scavenging system composed of PCA/PCD/Trolox). Purified motor protein was diluted to indicated concentrations in the assay buffer. Then, the solution was flowed into the glass chamber. An ECLIPSE Ti2-E microscope equipped with a CFI Apochromat TIRF 100XC Oil objective lens, an Andor iXion life 897 camera and a Ti2-LAPP illumination system (Nikon, Tokyo, Japan) was used to observe single molecule motility. NIS-Elements AR software ver. 5.2 (Nikon) was used to control the system.

## Acknowledgements

We would like to thank the members of the Niwa lab (Tohoku University). TK was supported by JSPS KAKENHI (23KJ0168). SN was supported by JSPS KAKENHI (23H02472 and 22H05523), the Naito foundation and the Uehara foundation. KC was supported by JSPS KAKENHI (22K15053) and the Naito foundation. Some worm strains and OP50 were obtained from the CGC.

## Statements

During the preparation of this manuscript, the authors used ChatGPT in order to check English grammar and improve English writing. After using this tool, the authors reviewed and edited the content as needed and take full responsibility for the content of the publication.

## Reference

Anazawa, Y., Kita, T., Iguchi, R., Hayashi, K. and Niwa, S. (2022). De novo mutations in KIF1A-associated neuronal disorder (KAND) dominant-negatively inhibit motor activity and axonal transport of synaptic vesicle precursors. Proc Natl Acad Sci U S A 119, e2113795119.

Arribere, J. A., Bell, R. T., Fu, B. X., Artiles, K. L., Hartman, P. S. and Fire, A. Z. (2014). Efficient marker-free recovery of custom genetic modifications with CRISPR/Cas9 in Caenorhabditis elegans. Genetics 198, 837–46.

Balseiro-Gomez, S., Park, J., Yue, Y., Ding, C., Shao, L., Etinkaya, S., Kuzoian, C., Hammarlund, M., Verhey, K. J. and Yogev, S. (2022). Neurexin and frizzled intercept axonal transport at microtubule minus ends to control synapse formation. Dev Cell 57, 1802–1816 e4.

Baron, D. M., Fenton, A. R., Saez-Atienzar, S., Giampetruzzi, A., Sreeram, A., Shankaracharya, Keagle, P. J., Doocy, V. R., Smith, N. J., Danielson, E. W., et al. (2022). ALS-associated KIF5A mutations abolish autoinhibition resulting in a toxic gain of function. Cell Rep 39, 110598.

Boyle, L., Rao, L., Kaur, S., Fan, X., Mebane, C., Hamm, L., Thornton, A., Ahrendsen, J. T., Anderson, M. P., Christodoulou, J. et al. (2021). Genotype and defects in microtubule-based motility correlate with clinical severity in KIF1A-associated neurological disorder. HGG Adv 2.

Brenner, S. (1974). The genetics of Caenorhabditis elegans. Genetics 77, 71–94.

Budaitis, B. G., Jariwala, S., Rao, L., Yue, Y., Sept, D., Verhey, K. J. and Gennerich, A. (2021). Pathogenic mutations in the kinesin-3 motor KIF1A diminish force generation and movement through allosteric mechanisms. J Cell Biol 220.

Chiba, K., Kita, T., Anazawa, Y. and Niwa, S. (2023). Insight into the regulation of axonal transport from the study of KIF1A-associated neurological disorder. J Cell Sci 136.

Chiba, K., Ori-McKenney, K. M., Niwa, S. and McKenney, R. J. (2022). Synergistic autoinhibition and activation mechanisms control kinesin-1 motor activity. Cell Rep 39, 110900.

Chiba, K., Takahashi, H., Chen, M., Obinata, H., Arai, S., Hashimoto, K., Oda, T., McKenney, R. J. and Niwa, S. (2019). Disease-associated mutations hyperactivate KIF1A motility and anterograde axonal transport of synaptic vesicle precursors. Proc Natl Acad Sci U S A 116, 18429–18434.

Ebbing, B., Mann, K., Starosta, A., Jaud, J., Schols, L., Schule, R. and Woehlke, G. (2008). Effect of spastic paraplegia mutations in KIF5A kinesin on transport activity. Hum Mol Genet 17, 1245–52.

Esmaeeli Nieh, S., Madou, M. R., Sirajuddin, M., Fregeau, B., McKnight, D., Lexa, K., Strober, J., Spaeth, C., Hallinan, B. E., Smaoui, N., et al. (2015). De novo mutations in KIF1A cause progressive encephalopathy and brain atrophy. Ann Clin Transl Neurol 2, 623–35.

Ghanta, K. S., Ishidate, T. and Mello, C. C. (2021). Microinjection for precision genome editing in Caenorhabditis elegans. STAR Protoc 2, 100748.

Glomb, O., Swaim, G., Munoz, L. P., Lovejoy, C., Sutradhar, S., Park, J., Wu, Y., Cason, S. E., Holzbaur, E. L. F., Hammarlund, M. et al. (2023). A kinesin-1 adaptor complex controls bimodal slow axonal transport of spectrin in Caenorhabditis elegans. Dev Cell 58, 1847–1863 e12.

Guardia, C. M., Farias, G. G., Jia, R., Pu, J. and Bonifacino, J. S. (2016). BORC Functions Upstream of Kinesins 1 and 3 to Coordinate Regional Movement of Lysosomes along Different Microtubule Tracks. Cell Rep 17, 1950–1961.

Hall, D. H. and Hedgecock, E. M. (1991). Kinesin-related gene unc-104 is required for axonal transport of synaptic vesicles in C. elegans. Cell 65, 837–47.

Hammond, J. W., Cai, D. W., Blasius, T. L., Li, Z., Jiang, Y. Y., Jih, G. T., Meyhofer, E. and Verhey, K. J. (2009). Mammalian Kinesin-3 Motors Are Dimeric In Vivo and Move by Processive Motility upon Release of Autoinhibition. Plos Biology 7, 650–663.

Higashida, M. and Niwa, S. (2023). Dynein intermediate chains DYCI-1 and WDR-60 have specific functions in Caenorhabditis elegans. Genes Cells 28, 97–110.

Hirokawa, N., Niwa, S. and Tanaka, Y. (2010). Molecular motors in neurons: transport mechanisms and roles in brain function, development, and disease. Neuron 68, 610–38.

Hirokawa, N., Noda, Y., Tanaka, Y. and Niwa, S. (2009). Kinesin superfamily motor proteins and intracellular transport. Nat Rev Mol Cell Biol 10, 682–96.

Holzbaur, E. L. and Scherer, S. S. (2011). Microtubules, axonal transport, and neuropathy. N Engl J Med 365, 2330–2.

Kita, T., Chiba, K., Wang, J., Nakagawa, A. and Niwa, S. (2023a). Comparative analysis of two Caenorhabditis elegans kinesins KLP-6 and UNC-104 reveals common and distinct activation mechanisms in kinesin-3. Elife 12, RP89040.

Kita, T., Sasaki, K. and Niwa, S. (2023b). Modeling the motion of disease-associated KIF1A heterodimers. Biophys J.

Klassen, M. P. and Shen, K. (2007). Wnt signaling positions neuromuscular connectivity by inhibiting synapse formation in C. elegans. Cell 130, 704–16.

Klassen, M. P., Wu, Y. E., Maeder, C. I., Nakae, I., Cueva, J. G., Lehrman, E. K., Tada, M., Gengyo-Ando, K., Wang, G. J., Goodman, M. et al. (2010). An Arf-like small G protein, ARL-8, promotes the axonal transport of presynaptic cargoes by suppressing vesicle aggregation. Neuron 66, 710-23.

Klebe, S., Lossos, A., Azzedine, H., Mundwiller, E., Sheffer, R., Gaussen, M., Marelli, C., Nawara, M., Carpentier, W., Meyer, V. et al. (2012). KIF1A missense mutations in SPG30, an autosomal recessive spastic paraplegia: distinct phenotypes according to the nature of the mutations. Eur J Hum Genet 20, 645–9.

Lam, A. J., Rao, L., Anazawa, Y., Okada, K., Chiba, K., Dacy, M., Niwa, S., Gennerich, A., Nowakowski, D. W. and McKenney, R. J. (2021). A highly conserved 310 helix within the kinesin motor domain is critical for kinesin function and human health. Sci Adv 7.

Li, S., Fell, S. M., Surova, O., Smedler, E., Wallis, K., Chen, Z. X., Hellman, U., Johnsen, J. I., Martinsson, T., Kenchappa, R. S. et al. (2016). The 1p36 Tumor Suppressor KIF 1Bbeta Is Required for Calcineurin Activation, Controlling Mitochondrial Fission and Apoptosis. Dev Cell 36, 164–78.

Luo, L. (2020). Principles of neurobiology, pp. 1 online resource. Boca Raton, FL: CRC Press, Taylor & Francis Group,.

Mello, C. C., Kramer, J. M., Stinchcomb, D. and Ambros, V. (1991). Efficient gene transfer in C.elegans: extrachromosomal maintenance and integration of transforming sequences. EMBO J 10, 3959–70.

Morikawa, M., Jerath, N. U., Ogawa, T., Morikawa, M., Tanaka, Y., Shy, M. E., Zuchner, S. and Hirokawa, N. (2022). A neuropathy-associated kinesin KIF1A mutation hyper-stabilizes the motor-neck interaction during the ATPase cycle. EMBO J 41, e108899.

Munirajan, A. K., Ando, K., Mukai, A., Takahashi, M., Suenaga, Y., Ohira, M., Koda, T., Hirota, T., Ozaki, T. and Nakagawara, A. (2008). KIF1Bbeta functions as a haploinsufficient tumor suppressor gene mapped to chromosome 1p36.2 by inducing apoptotic cell death. J Biol Chem 283, 24426–34.

Nakano, J., Chiba, K. and Niwa, S. (2022). An ALS-associated KIF5A mutant forms oligomers and aggregates and induces neuronal toxicity. Genes Cells 27, 421–435.

Nangaku, M., Sato-Yoshitake, R., Okada, Y., Noda, Y., Takemura, R., Yamazaki, H. and Hirokawa, N. (1994). KIF1B, a novel microtubule plus end-directed monomeric motor protein for transport of mitochondria. Cell 79, 1209–20.

Niwa, S., Lipton, D. M., Morikawa, M., Zhao, C., Hirokawa, N., Lu, H. and Shen, K. (2016). Autoinhibition of a Neuronal Kinesin UNC-104/KIF1A Regulates the Size and Density of Synapses. Cell Rep 16, 2129–41.

Niwa, S., Tanaka, Y. and Hirokawa, N. (2008). KIF1Bbeta-and KIF1A-mediated axonal transport of presynaptic regulator Rab3 occurs in a GTP-dependent manner through DENN/MADD. Nat Cell Biol 10, 1269–79.

Okada, Y., Yamazaki, H., Sekine-Aizawa, Y. and Hirokawa, N. (1995). The neuron-specific kinesin superfamily protein KIF1A is a unique monomeric motor for anterograde axonal transport of synaptic vesicle precursors. Cell 81, 769–80.

Otsuka, A. J., Jeyaprakash, A., Garcia-Anoveros, J., Tang, L. Z., Fisk, G., Hartshorne, T., Franco, R. and Born, T. (1991). The C. elegans unc-104 gene encodes a putative kinesin heavy chain-like protein. Neuron 6, 113–22.

Pack-Chung, E., Kurshan, P. T., Dickman, D. K. and Schwarz, T. L. (2007). A Drosophila kinesin required for synaptic bouton formation and synaptic vesicle transport. Nat Neurosci 10, 980–9.

Pant, D. C., Parameswaran, J., Rao, L., Loss, I., Chilukuri, G., Parlato, R., Shi, L., Glass, J. D., Bassell, G. J., Koch, P. et al. (2022). ALS-linked KIF5A DeltaExon27 mutant causes neuronal toxicity through gain-of-function. EMBO Rep 23, e54234.

Pierce-Shimomura, J. T., Chen, B. L., Mun, J. J., Ho, R., Sarkis, R. and McIntire, S. L. (2008). Genetic analysis of crawling and swimming locomotory patterns in C. elegans. Proc Natl Acad Sci U S A 105, 20982–7.

Ren, J., Wang, S., Chen, H., Wang, W., Huo, L. and Feng, W. (2018). Coiled-coil 1-mediated fastening of the neck and motor domains for kinesin-3 autoinhibition. Proc Natl Acad Sci U S A 115, E11933–E11942.

Stucchi, R., Plucinska, G., Hummel, J. J. A., Zahavi, E. E., Guerra San Juan, I., Klykov, O., Scheltema, R. A., Altelaar, A. F. M. and Hoogenraad, C. C. (2018). Regulation of KIF1A-Driven Dense Core Vesicle Transport: Ca(2+)/CaM Controls DCV Binding and Liprin-alpha/TANC2 Recruits DCVs to Postsynaptic Sites. Cell Rep 24, 685–700.

Tan, Z., Yue, Y., da Veiga Leprevost, F., Haynes, S. E., Basrur, V., Nesvizhskii, A. I., Verhey, K. J. and Cianfrocco, M. A. (2023). Autoinhibited kinesin-1 adopts a hierarchical folding pattern. Elife 12, RP86776.

Tomishige, M., Klopfenstein, D. R. and Vale, R. D. (2002). Conversion of Unc104/KIF1A kinesin into a processive motor after dimerization. Science 297, 2263–2267.

Verhey, K. J. and Hammond, J. W. (2009). Traffic control: regulation of kinesin motors. Nature Reviews Molecular Cell Biology 10, 765–777.

Wang, W., Ren, J., Song, W., Zhang, Y. and Feng, W. (2022). The architecture of kinesin-3 KLP-6 reveals a multilevel-lockdown mechanism for autoinhibition. Nat Commun 13, 4281.

Weijman, J. F., Yadav, S. K. N., Surridge, K. J., Cross, J. A., Borucu, U., Mantell, J., Woolfson, D. N., Schaffitzel, C. and Dodding, M. P. (2022). Molecular architecture of the autoinhibited kinesin-1 lambda particle. Sci Adv 8, eabp9660.

Wu, Y. E., Huo, L., Maeder, C. I., Feng, W. and Shen, K. (2013). The balance between capture and dissociation of presynaptic proteins controls the spatial distribution of synapses. Neuron 78, 994–1011.

Xu, F., Takahashi, H., Tanaka, Y., Ichinose, S., Niwa, S., Wicklund, M. P. and Hirokawa, N. (2018). KIF1Bβ mutations detected in hereditary neuropathy impair IGF1R transport and axon growth. Journal of Cell Biology 217, 3480–3496.

Zhao, C., Takita, J., Tanaka, Y., Setou, M., Nakagawa, T., Takeda, S., Yang, H. W., Terada, S., Nakata, T., Takei, Y. et al. (2001). Charcot-Marie-Tooth disease type 2A caused by mutation in a microtubule motor KIF1Bbeta. Cell 105, 587–97.

Zuchner, S., Mersiyanova, I. V., Muglia, M., Bissar-Tadmouri, N., Rochelle, J., Dadali, E. L., Zappia, M., Nelis, E., Patitucci, A., Senderek, J. et al. (2004). Mutations in the mitochondrial GTPase mitofusin 2 cause Charcot-Marie-Tooth neuropathy type 2A. Nat Genet 36, 449–51.

