## Supplemental for "Characterizing Human KIF1Bß Motor Activity by Single-Molecule Motility Assays and *Caenorhabtidis elegans* Genetics"

| Plasmid name | insert | comment |
|---|---|---|
| pSN721 | Punc-104::KIF1Bbeta | Fig 1 and 2 |
| pSN993 | Punc-104::KIF1Bbeta (Q98L) | Fig 1 and 2 |
| pSN541 | pAcebac1_hsKIF1Bbeta(codon optimized)_sfGFP::2xStrep tag | Fig3 |
| pSN667 | pAcebac1_hKIF1Bbeta(1-721)(codon optimized)_sfGFP_strepII | Fig 3 and 4 |
| pSN773 | pAcebac1_hKIF1Bbeta(1-721)(Q98L)(codon optimized)_sfGFP_strepII | Fig 3 and 4 |

#### Supplementary Table S1 Plasmid List

| Strain Name | Genotype | Comment |
|---|---|---|
| N2 | *wild type* | |
| TV1229 | *wyls85* V | |
| OTL129 | *unc-104(e1265)* II; *wyls85* V | |
| OTL235 | *unc-104(e1265)* ; *wyls85* ; *jpnEx571* [Punc-104::Kif1bß ] | Figure 1and Figure 2 |
| OTL236 | *unc-104(e1265)* ; *wyls85* ; *jpnEx572* [Punc-104::Kif1bß(Q98L) ] | Figure 1and Figure 2 strain #1 |
| OTL237 | *unc-104(e1265)* ; *wyls85* ; *jpnEx573* [Punc-104::Kif1bß(Q98L) ] | Figure 1and Figure 2 strain #2 |
| OTL201 | *unc-104(jpn61)* II; *wyls85* V | jpn61 is unc-104(Q94L) mutant. Figure 5 and Figure 6 |

### Supplementary Table S2 Strain List

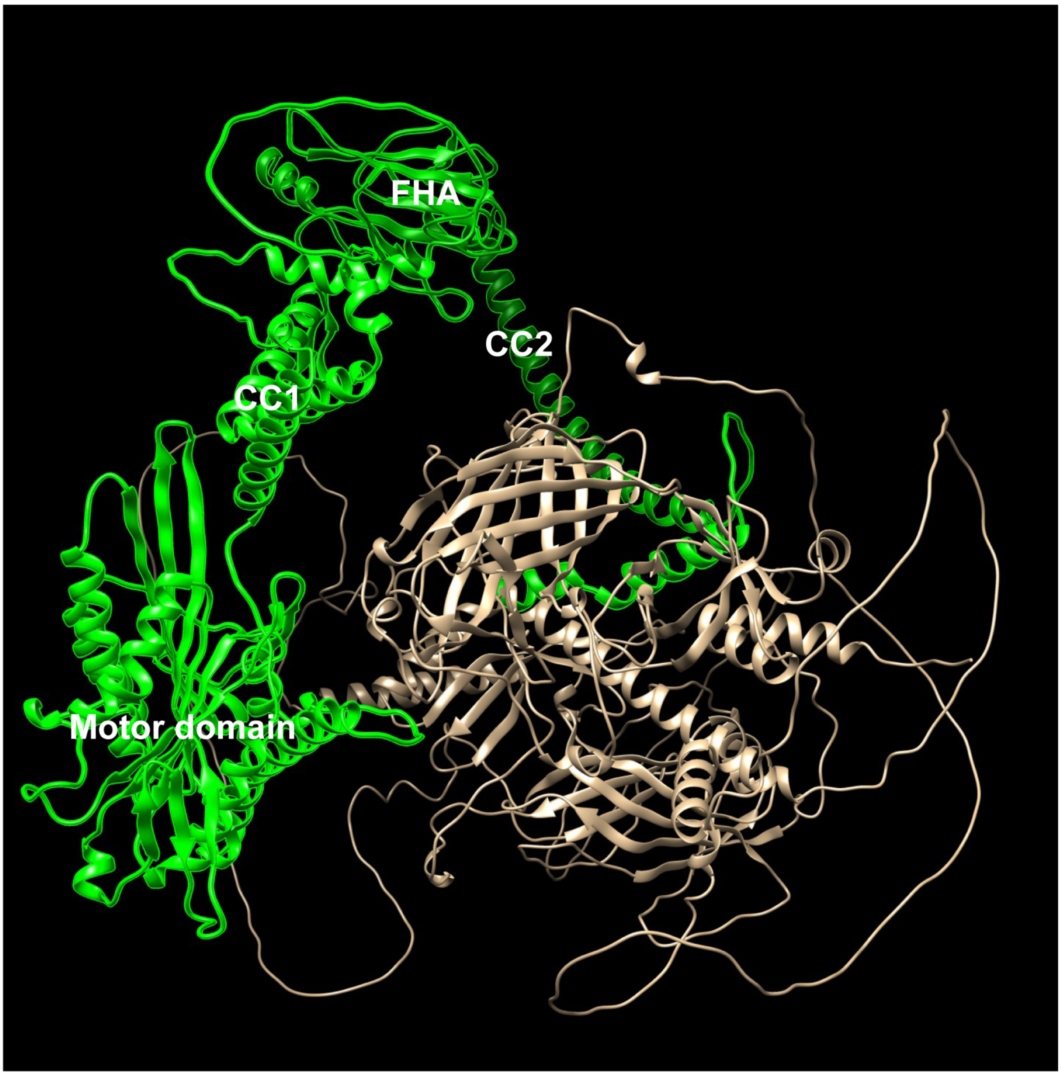


**Figure S1 Predicted KIF1Bß structure**

Human KIF1Bß was obtained from AlphaFold Protein Structure Database (Varadi et al., 2022). KIF1Bß(1-721) is shown by green.

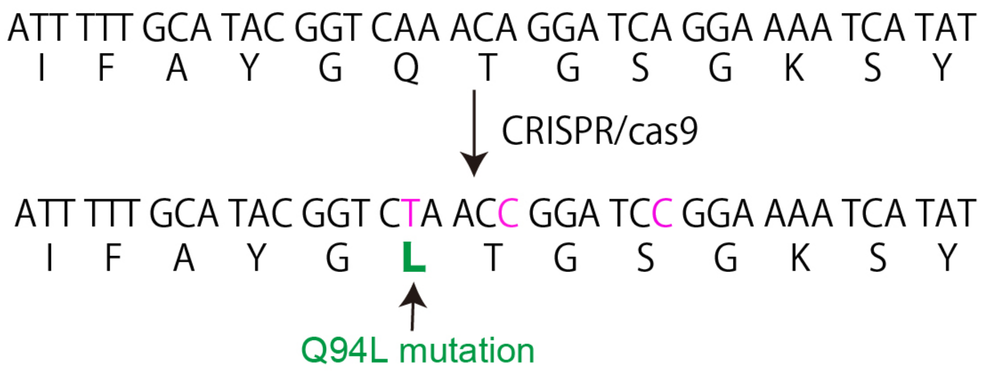


**Figure S2 Schematic drawing showing the genome editing to introduce Q94L mutation**

The genome sequence of *unc-104* encoding UNC-104(Q94) (upper) and edited sequence (lower) are shown.

**Reference**

**Varadi, M., Anyango, S., Deshpande, M., Nair, S., Natassia, C., Yordanova, G., Yuan, D., Stroe, O., Wood, G., Laydon, A. et al.** (2022). AlphaFold Protein Structure Database: massively expanding the structural coverage of protein-sequence space with high-accuracy models. *Nucleic Acids Res* **50**, D439-D444.
